# Inhibition of immunoglobulin class-switching prevents pemphigus onset in desmoglein 3-specific B cell receptor knock-in mouse

**DOI:** 10.1101/2021.05.23.445300

**Authors:** Hisashi Nomura, Naoko Wada, Hayato Takahashi, Yuko Kase, Jun Yamagami, Shohei Egami, Hisato Iriki, Miho Mukai, Aki Kamata, Hiromi Ito, Hideki Fujii, Tomoyuki Ishikura, Haruhiko Koseki, Takashi Watanabe, Taketo Yamada, Osamu Ohara, Shigeo Koyasu, Masayuki Amagai

## Abstract

Although immunoglobulin class-switching is essential for humoral immunity, its role in B-cell immune tolerance remains unclear. Pemphigus vulgaris is an autoimmune blistering disease caused by IgG targeting desmoglein 3, an adhesion molecule of keratinocytes. In this study, we generated knock-in mice that express anti-Dsg3 AK23 autoantibodies. Knock-in B cells developed normally *in vivo* and showed Ca^2+^ influx upon IgM cross-linking *in vitro*. The mice predominantly produced circulating AK23 IgM but little IgG antibodies. Although no IgG deposition or blister formation was observed in Dsg3-bearing tissues, Dsg3 immunization forced to induce pemphigus phenotype after class-switching to IgG *in vivo*. Transcriptomic analysis revealed that *FCGR2B* and FcγRIIB-related genes were downregulated in B cells from peripheral blood of pemphigus patients. Indeed, in AK23 knock-in mice, *Fcgr2b* deficiency or haploinsufficiency spontaneously led to class-switching, AK23 IgG production, and pemphigus phenotype development. Thus, inhibition of pathogenic class-switching is a crucial tolerogenic process to prevent pemphigus onset, where attenuated FcγRIIB signaling is one of the key predispositions to break this tolerogenic state.

## INTRODUCTION

Immunological tolerance is crucial in preventing tissue injury caused by autoreactive lymphocytes. Clonal deletion (1), anergy, (2) and receptor editing (3, 4) are widely accepted B cell tolerance mechanisms. Since tolerance is achieved during key processes that are necessary for B cells to develop and function properly, harmful activities of autoreactive B cells can be efficiently constrained. In anergy, for example, activation of autoreactive B cells after antigen recognition by B cell receptor (BCR) is inhibited (2, 5, 6), while immunoglobulin autoreactivity can be reversed via antibody gene rearrangement, also referred to as receptor editing (3, 4, 7). The most basic function of B cells is efficient antibody production, and its essential steps include somatic hypermutation, class-switching of immunoglobulin, differentiation into memory B cells and antibody-producing plasma cells and so on (8–10). The tolerogenic processes involved in these steps must be efficient to prevent autoimmunity. Indeed, the limited accumulation of IgG^+^ anti-DNA plasma cells was reported as a crucial process to prevent autoantibody-mediated glomerulonephritis (11). Nevertheless, tolerogenic actions to the most steps necessary for antibody production have not been elucidated.

Pemphigus vulgaris (PV) is a life-threatening autoimmune blistering disease induced by IgG against desmoglein 3 (Dsg3), a cadherin-type cell adhesion molecule expressed in the stratified squamous epithelium (12, 13). Since IgG interferes with cell-cell adhesion, keratinocytes become dissociated, resulting in blister formation in the skin and oral mucosa. Anti-Dsg3 IgG is detected in PV patients but not in healthy individuals (14); however, the mechanisms underlying autoantibody production are not fully understood. PV mouse model was previously established by immunizing *Dsg3*^-/-^ mice with recombinant Dsg3 (rDsg3); since *Dsg3*^-/-^ mice lack tolerance against Dsg3, immunization with rDsg3 results in potent Dsg3-specific immune reactions (15). Upon adoptive transfer of lymphocytes from immunized *Dsg3*^-/-^ mice to *Rag2*^-/-^ mice, Dsg3-specific lymphocytes expand and produce anti-Dsg3 IgG, resulting in PV phenotype. A series of anti-Dsg3 antibody clones were isolated from this model for further characterization and epitope mapping (16).

AK23 is a mouse IgG_1_ antibody clone that recognizes Dsg3 and causes blisters *in vivo* To identify novel tolerance mechanisms in Dsg3-specific B cells, AK23 IgM transgenic mice were generated (17). These mice have AK23 heavy (H) and light (L) chains in B cells, which produce IgM with AK23 antigen-specificity. Although AK23 IgM-expressing B cells and circulating AK23 IgM were detected in transgenic mice (17), no profound IgM deposition on the palate was observed, and the mice did not develop PV phenotype. Electron microscopy revealed AK23 IgM deposition primarily at the periphery of desmosomes, whereas AK23 IgG accumulated at the center of desmosomes (17). These results implied that AK23 IgM was non-pathogenic *in vivo* because it could not access Dsg3 at the desmosome core, likely due to the larger size of IgM compared to IgG. Since IgM transgenic mice are incapable of immunoglobulin class-switching, accurate investigation on tolerance mechanisms against pathogenic autoreactive B cells required BCR knock-in mice that allow immunoglobulin class-switching *in vivo*.

In this study, we generated Dsg3-specific AK23 BCR knock-in mouse to explore tolerance mechanisms in autoreactive B cells. Downregulation of FcγRIIB signaling, a pathway often dysregulated in B cells of PV patients, promoted PV phenotype in mice by driving “pathogenic” immunoglobulin class-switching. Our results unveiled a crucial tolerogenic process that acts by modulating immunoglobulin class-switching in B cells.

## RESULTS

### Dsg3-specific BCR knock-in B cells exhibit attenuated pathogenicity *in vivo*

To generate a Dsg3-specific BCR knock-in mouse line, we constructed an immunoglobulin H chain knock-in vector, wherein the endogenous J_H_ gene cluster was replaced with the VDJ gene segment of the AK23 H chain gene (**Fig. 1A**). AK23 H chain-knock-in mice (AK23H^ki^) were generated by injecting the knock-in vector into v6.5 mouse embryonic stem (ES) cells (C57BL/6x129/Sv) (**Fig. 1B**); AK23H^ki^ mice were backcrossed to C57BL/6 mice for more than ten generations. AK23H^ki^ mice were then crossed with AK23 L chain transgenic mice (17) to generate AK23 knock-in mice (AK23H^ki^L^tg^). To confirm Dsg3-reactivity of B cells generated in AK23H^ki^L^tg^ mice, we performed flow cytometry after incubation of the B cells with recombinant Dsg3 protein. As expected, Dsg3-bound B cells were detected as a single population positively shifted in plots (**Fig. 1C**). Indeed, when we analyzed whether the targeted allele harbored IgM^a^ or IgM^b^ allotypes originating from 129/Sv or C57BL/6 mice, respectively, we found that B cells isolated from AK23H^ki^ mice expressed the IgM^a^, but not IgM^b^, allotype (**Fig. 1D**). These findings imply that Dsg3-specific AK23 BCR was successfully expressed in B cells as IgM^a^ due to allelic exclusion in AK23H^ki^L^tg^ mice.

**Figure 1.**
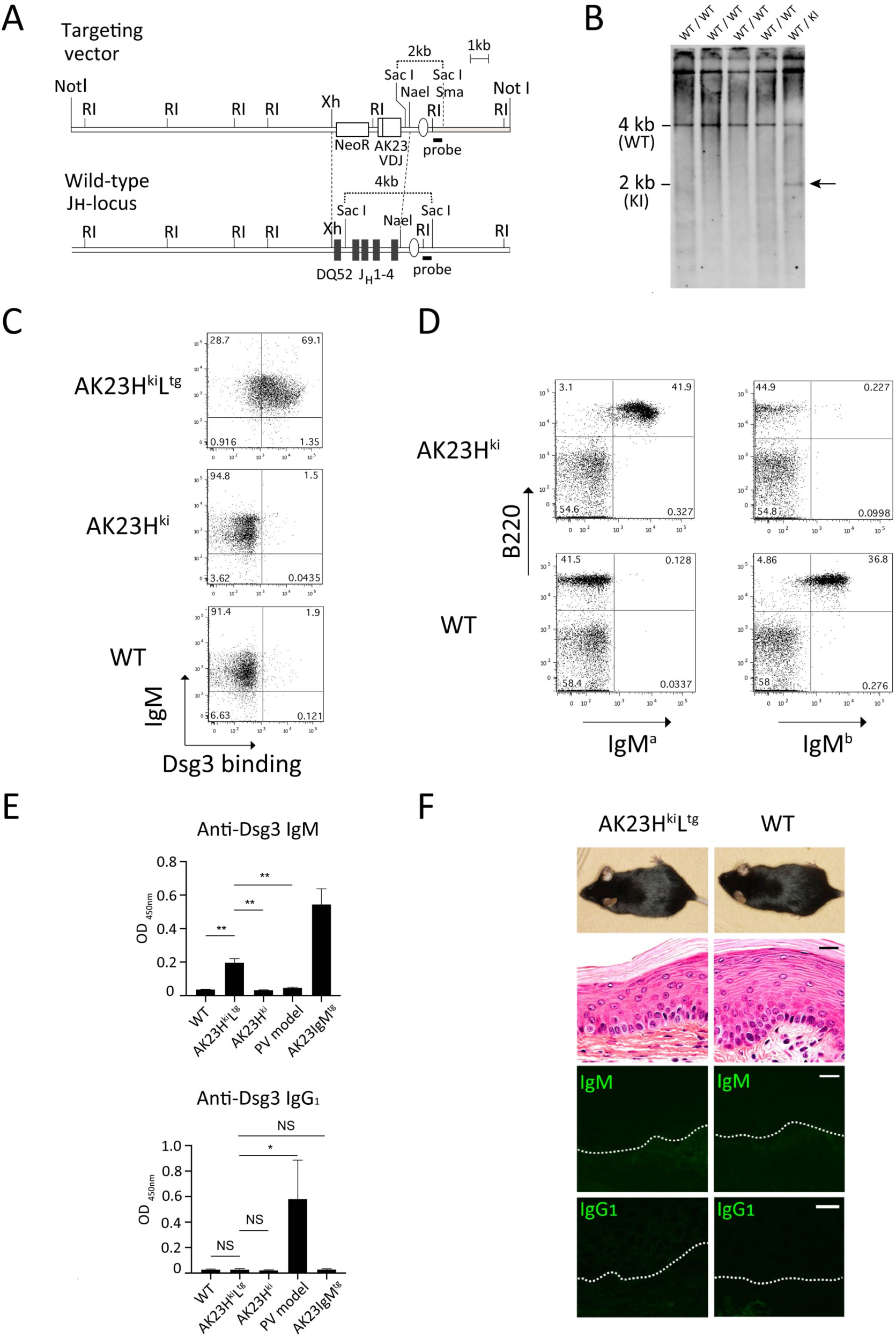
Dsg3-specific BCR knock-in B cells exhibit attenuated pathogenicity *in vivo*. (**A**) Scheme of the AK23 heavy chain knock-in vector. The probe used for Southern blot analysis of **Fig. 1B** is indicated with a thick black line. (**B**) Southern blot analysis of knocked-in ES cells. The WT and knocked-in alleles were detected as 4 kb and 2 kb fragments, respectively. (**C**) Dsg3 specificities of splenocytes from AK23H^ki^L^tg^ (n = 2), AK23H^ki^ (n = 2), and WT (n = 2) mice by flow cytometry. (**D**) Splenocytes from AK23H^ki^ (n = 2) and WT (n = 2) mice determined by flow cytometry. (**E**) Serum anti-Dsg3 IgM and IgG_1_ levels in WT (n = 6), AK23H^ki^ (n = 7), and AK23H^ki^L^tg^ (n = 6) mice were determined by ELISA. Serum from AK23IgM^tg^ mice (n = 3) and PV model mice (n = 3) were used as a positive control for anti-Dsg3 IgM and IgG_1_ quantification, respectively. (**F**) Skin phenotype, histopathology, and immunofluorescence in AK23H^ki^L^tg^ and WT mice. Dashed lines indicate basement membrane zones. Scale bar, 20 μm. Data are shown as the mean ± SEM. **Figure 1-Source Data 1** The original figure file of the full raw unedited gel shown in **Fig. 1B**. **Figure 1-Source Data 2** The file of the figure with the uncropped gel with the relevant bands labelled in **Fig. 1B**. **Figure 1-Source Data 3** Excel file containing numerical values collected from ELISA of anti-Dsg3 IgM and IgG_1_ titer shown in **Fig. 1E**.

Next, we assessed tissue deposition of circulating immunoglobulins. In contrast to AK23H^ki^ or wild-type (WT) mice, circulating anti-Dsg3 IgM was abundant in AK23H^ki^L^tg^ mice (**Fig. 1E**). However, immunofluorescence staining revealed that *in vivo* IgM deposition on keratinocyte cell surfaces occurred at extremely low levels in the palate of AK23H^ki^L^tg^ mice (**Fig. 1F**). Although Dsg3-specific AK23 BCR knock-in B cells maintained the potential for class-switching, only marginal levels of circulating anti-Dsg3 IgG1 were detected in AK23H^ki^L^tg^ mice, which were considerably lower than in PV model mice (**Fig. 1E**). Consistently, *in vivo* IgG1 deposition in the palate was minimal in AK23H^ki^L^tg^ mice (**Fig. 1F**). In line with these results, we did not observe skin erosion, hair loss, acantholysis, or other pemphigus phenotype in AK23H^ki^L^tg^ mice (**Fig. 1F**). Furthermore, up to 1 year of age, neither serum anti-Dsg3 IgG1 nor *in vivo* IgG1 deposition levels spontaneously increased to the levels that induce acantholytic blister. These results imply that despite the Dsg3 reactivity of AK23H^ki^L^tg^ B cells, their pathogenicity was attenuated, and differentiation into pathogenic IgG^+^ B cells was not observed even in the presence of Dsg3.

### AK23H^ki^L^tg^ B cells are capable of BCR-mediated activation but remain non-pathogenic at a steady state *in vivo*

Previous studies have shown that partial clonal deletion and anergy occurred even in BCR transgenic mice that produced autoreactive IgM antibodies (18). Although most AK23H^ki^L^tg^ B cells expressed IgM, AK23H^ki^L^tg^ B cells may undergo clonal deletion or anergy. Thus, we assessed AK23H^ki^L^tg^ B cell deletion or anergy in the presence and absence of Dsg3. To this end, we transplanted bone marrow (BM) cells from AK23H^ki^L^tg^-*Rag2^-/-^* mice (CD45.1^+^) into irradiated WT and *Dsg3*^-/-^ mice (CD45.2^+^) (**Fig. 2A**) and assessed AK23H^ki^L^tg^ B cell development in the presence and absence of Dsg3, respectively. Hardy fraction, a representative classification, was utilized for BM analysis (19). We found no significant difference in the ratio of Hardy fractions A to F during AK23H^ki^L^tg^ B cell development in the two groups (**Fig. 2B**). We performed similar analyses using splenic B cells, which consist of cells at transitional stages (T1, T2, and T3), follicular B cells, and marginal zone B cells (20). No significant differences in these subpopulations between the two groups were observed (**Fig. 2C**). These findings imply the lack of Dsg3-dependent clonal deletion in AK23H^ki^L^tg^ B cells.

**Figure 2.**
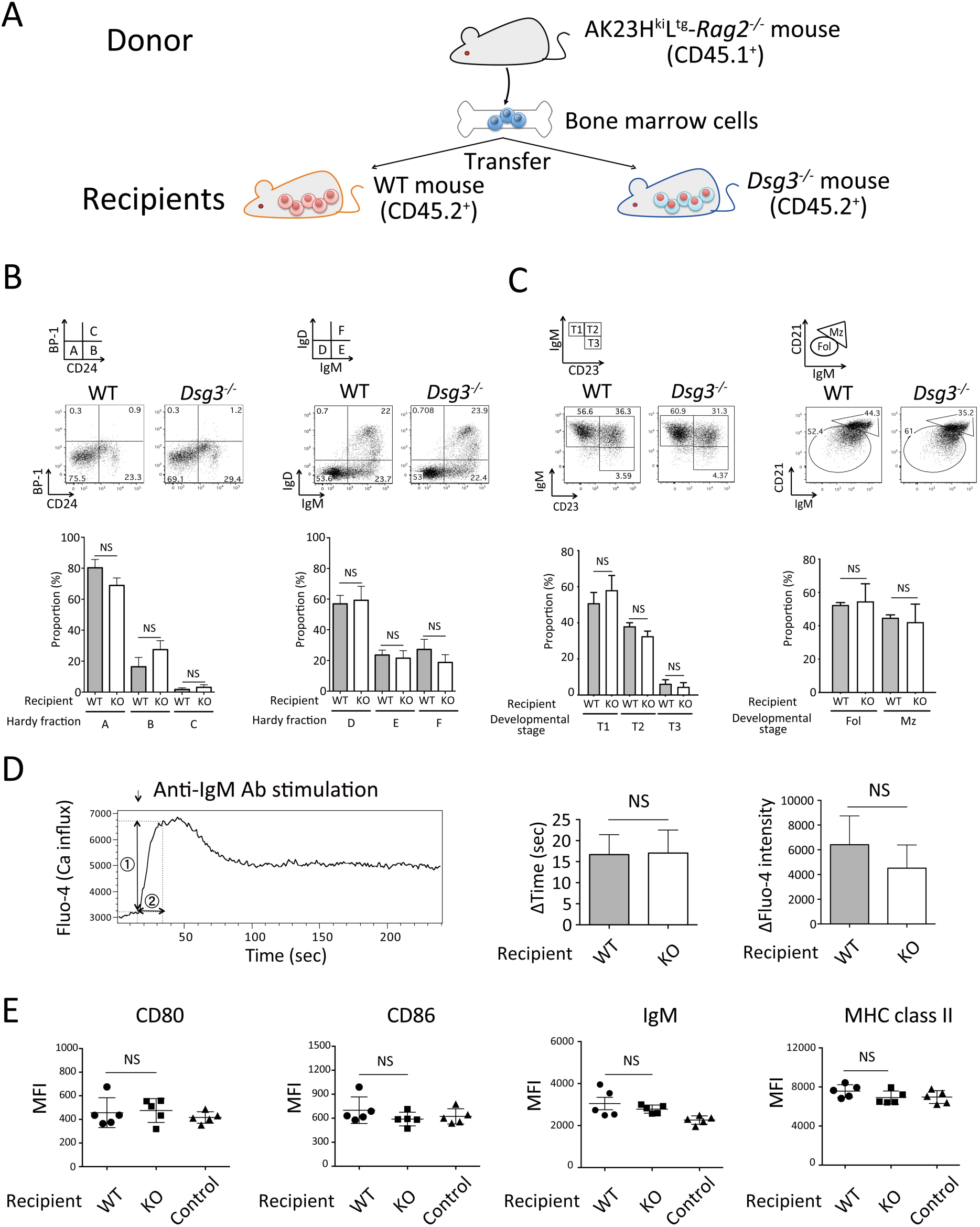
AK23H^ki^L^tg^ B cells are not subjected to clonal deletion or anergy. (**A**) Scheme of bone marrow transfer (BMT) experiments. All experiments were performed with AK23H^ki^L^tg^ B cells from WT and *Dsg3^-/-^* recipients after BMT. (**B**) AK23H^ki^L^tg^ B cell development in WT and *Dsg3^-/-^* recipients assessed by flow cytometry. Hardy fraction A–C and D–F B cells were identified after gating on CD45.1^+^B220^+^CD43^high^ and CD45.1^+^B220^+^CD43^low^ cells, respectively (**Fig. S3A**). Representative flow cytometry plots, and quantitative summaries on proportions of each developmental stage (n = 3) are shown. (**C**) Splenic AK23H^ki^L^tg^ B cell development in WT and *Dsg3^-/-^* recipients assessed by flow cytometry. Donor-derived transitional (T1, T2, and T3) and follicular (Fol)/marginal zone (Mz) B cells were identified after gating on CD45.1^+^Gr-1^-^CD11b^-^CD19^+^B220^high^CD93^+^ and CD45.1^+^Gr-1^-^CD11b^-^CD19^+^B220^high^CD93^-^ cells, respectively (**Fig. S3B**). Proportions of these B cells are shown in representative flow cytometry plots and bar graphs (n = 3). (**D**) Ca influx in splenic AK23H^ki^L^tg^ B cells from WT and *Dsg3^-/-^* recipients after anti-IgM antibody stimulation (arrow) assessed by flow cytometry. Fluorescence intensity (1) and the time (2) between anti-IgM stimulation and fluorescence intensity peak were determined (n = 4). (**E**) Quantitation of CD80, CD86, surface IgM, and MHC class II expressions examined by flow cytometry in splenic AK23H^ki^L^tg^ B cells from WT and *Dsg3^-/-^* recipients. WT B cells from WT recipients after BMT from WT mice were used as controls (n = 5). MFI, mean fluorescence intensity. Data are shown as the mean ± SEM. **Figure 2-Source Data 1** Excel file containing numerical values of the analyses shown in **Fig. 2B-E**.

To assess whether AK23H^ki^L^tg^ B cells are subjected to anergy, we isolated AK23H^ki^L^tg^ B cells from the BM of chimeric recipients (**Fig. 2A**) and analyzed their responses to anti-IgM stimulation using Fluo-4, a dye that emits fluorescence upon calcium influx after antigen stimulation. When AK23H^ki^L^tg^ B cells derived from WT mice were stimulated, calcium influx was observed by flow cytometry (**Fig. 2D**). To statistically analyze the difference in antigen-initiated B cell reactivity, we considered the difference in fluorescence intensity and the time between stimulation and peak fluorescence intensity; no significant difference was detected (**Fig. 2D**). In addition, expression levels of cell surface markers such as CD80, CD86, IgM, and MHC class II, which are usually attenuated in the anergic state, were also comparable between these two groups (**Fig. 2E**), implying no differences in the levels of B cell anergy and that AK23H^ki^L^tg^ B cells were not subjected to clonal deletion or anergy. Instead, AK23H^ki^L^tg^ B cells remained non-pathogenic in a steady state in secondary lymphoid organs, retaining their potential to respond to BCR stimulation.

### AK23H^ki^L^tg^ B cells show pathogenic potential for PV phenotype induction after *in vitro* class-switching

To investigate whether AK23H^ki^L^tg^ B cells produce IgG autoantibodies after class-switching, we forced class-switching to IgG_1_ by stimulating B cells with IL-4 and LPS *in vitro* (**Fig. 3A**). We found that an AK23H^ki^L^tg^ B cell subpopulation was converted into IgG_1_^+^ B cells, and most of these cells were reactive to rDsg3 protein (**Fig. 3B**). After adoptive transfer of IgG_1_-producing AK23H^ki^L^tg^ B cells into *Rag2^-/-^* mice, transplanted mice developed skin erosions, crusted skin lesions, and hair loss in the skin (**Fig. 3C**). Furthermore, immunofluorescence analysis revealed IgG_1_ deposition on keratinocyte cell surfaces and acantholytic blister, the two most characteristic features of pemphigus (**Fig. 3C**). These findings indicate that AK23H^ki^L^tg^ B cells retained their potential for class-switching *in vitro* and that IgG autoantibodies produced by AK23H^ki^L^tg^ B cells were pathogenic *in vivo*.

**Figure 3.**
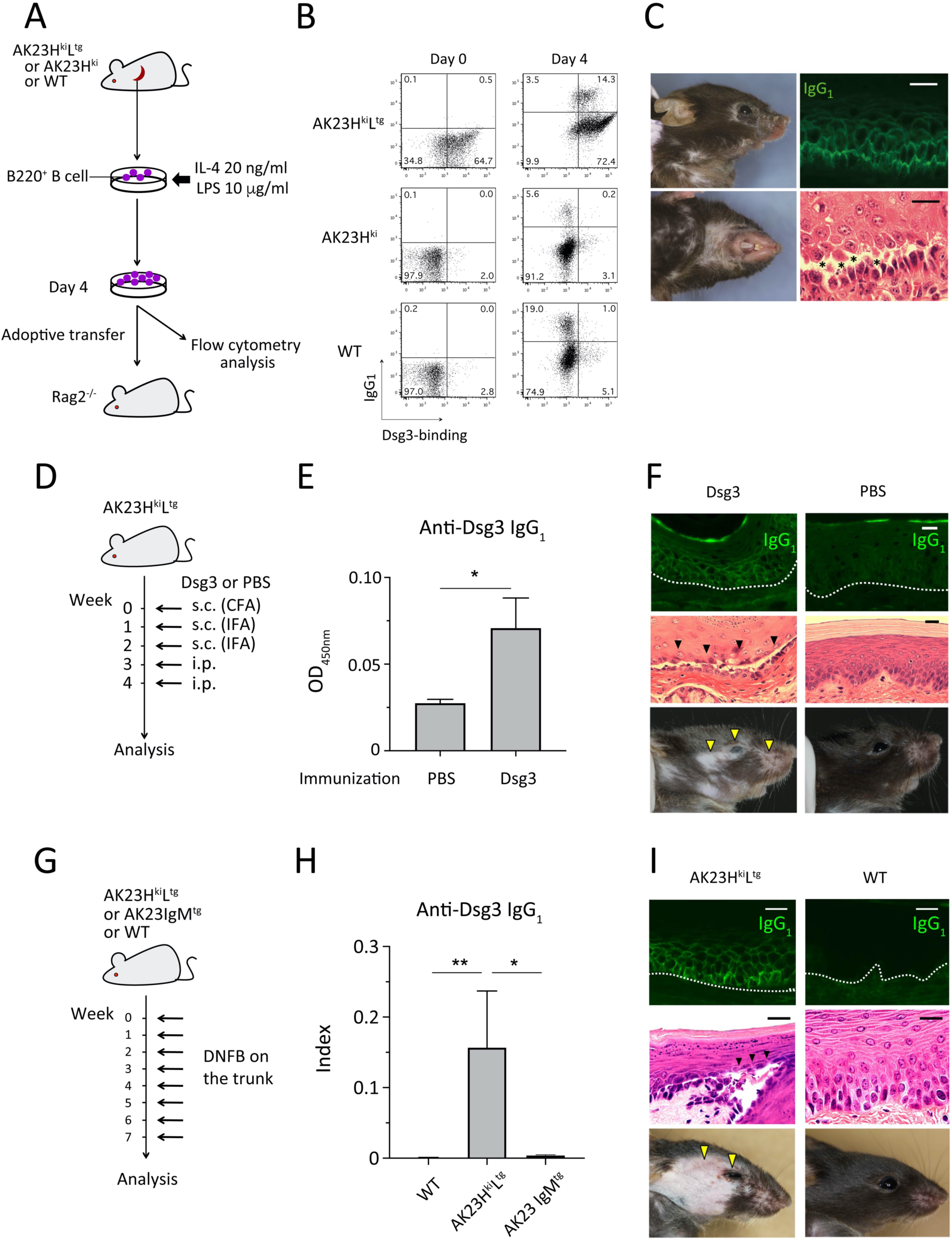
Immunoglobulin class-switching in AK23H^ki^L^tg^ B cells initiates PV phenotype induction. (**A**) Schematic outline of the experimental procedures. (**B**) Flow cytometry analysis of splenic B cells from AK23H^ki^L^tg^, AK23H^ki^, and WT mice before (day 0) and after (day 4) *in vitro* stimulation with IL-4 and LPS to detect Dsg3-specific IgG_1_^+^ B cells. (**C**) Clinical presentation, hematoxylin and eosin staining, and immunofluorescence staining of *Rag2^-/-^* recipients (n = 2) after adoptive transfer of AK23H^ki^L^tg^ IgG_1_^+^ B cells. PV phenotypes, including skin erosions and hair loss, were recorded. IgG_1_ deposition on keratinocyte cell surfaces (green) and acantholysis (*) are shown. Scale bar, 20 μm. (**D**) Scheme of immunization procedure. S.c., subcutaneous injection; i.p., intraperitoneal injection; CFA, complete Freund’s adjuvant; IFA, incomplete Freund’s adjuvant. (**E**) Serum anti-Dsg3 IgG_1_ were quantified by ELISA in AK23H^ki^L^tg^ mice after serial immunization with rDsg3 (n = 5) and PBS (n = 4). Data are pooled from 2 independent experiments. (**F**) Skin phenotype of Dsg3-immunized AK23H^ki^L^tg^ and control mice. IgG_1_ depositions shown in green, acantholytic blister indicated by black arrowheads, and hair loss and skin erosion indicated by yellow arrowheads. (**G**) Scheme of DNFB treatment. DNFB was weekly applied to the trunk of AK23H^ki^L^tg^, WT and AK23IgM^tg^ mice (**H**) Anti-Dsg3 IgG_1_ antibody were quantified after DNFB treatment of AK23H^ki^L^tg^ (n = 8), WT (n = 4) and AK23IgM^tg^ (n = 3) mice by ELISA. Data are pooled from 2 independent experiments. (**I**) Skin phenotype of AK23H^ki^L^tg^ mice after DNFB treatment. Hair loss and erosion indicated by yellow arrowheads, acantholytic blister indicated by black arrowheads, and IgG_1_ depositions shown in green. Scale bar, 20 μm. \**P* < 0.05, \*\**P* < 0.01. Statistical significance was determined by Mann-Whitney *U* test (E, H). Data are shown as the mean ± SEM. **Figure 3-Source Data 1** Excel file containing numerical values collected from ELISA of anti-Dsg3 IgG_1_ titer shown in **Fig. 3E and H.**

### Forced class-switching in Dsg3-specific BCR knock-in B cells by Dsg3 immunization and skin damage induces the PV phenotype *in vivo*

To assess whether class-switching in AK23H^ki^L^tg^ B cells can be induced *in vivo*, we repeatedly immunized AK23H^ki^L^tg^ mice with rDsg3 protein and adjuvant (**Fig. 3D**). In contrast to mice treated with adjuvant alone, rDsg3-immunized AK23H^ki^L^tg^ mice started producing anti-Dsg3 IgG_1_ antibodies and developed a PV phenotype at approximately 4 weeks after treatment (**Fig. 3E, 3F**). Consistently, acantholytic blister and IgG_1_ deposition on keratinocyte cell surfaces were histologically detected in rDsg3-immunized AK23H^ki^L^tg^ mice but not in mice immunized with adjuvant alone (**Fig. 3F**), indicating that Dsg3 immunization promoted class-switching in Dsg3-specific BCR knock-in B cells *in vivo*.

To evaluate whether endogenous Dsg3 exposure of the immune system is sufficient to drive class-switching in AK23H^ki^L^tg^ B cells, we employed a hapten-induced dermatitis model, where Dsg3-expressing keratinocytes were damaged by forced dermatitis. Repeated application of 1-fluoro-2,4-dinitrobenzene (DNFB) on shaved trunk skin induced severe dermatitis and skin erosion (**Fig. 3G**). Additionally, DNFB treatment promoted anti-Dsg3 IgG_1_ production in AK23H^ki^L^tg^ mice but not in WT or AK23IgM^tg^ mice (**Fig. 3H**). In fact, the PV phenotype including erosions and hair loss was observed on the face of AK23H^ki^L^tg^ mice, on which DNFB was not applied (**Fig. 3I**). On the other hand, dermatitis did not develop on the face in WT and AK23IgM^tg^ mice (**Fig. 3I**, and data not shown). In addition, acantholytic blister and IgG_1_ deposition on the keratinocyte cell surfaces were confirmed in AK23H^ki^L^tg^ mice (**Fig. 3I**).

Viral infections have been implicated in autoimmune disease exacerbation (21). Hence, we assessed the ability of a skin-tropic virus, vaccinia virus, to induce class-switching in AK23H^ki^L^tg^ B cells by damaging Dsg3-bearing keratinocytes. To this end, we inoculated the trunk skin of AK23H^ki^L^tg^ mice with vaccinia virus (**Fig. S1A**). After repeated inoculation, we observed skin erosion and IgG_1_ deposition on keratinocyte cell surfaces of AK23H^ki^L^tg^ mice (**Fig. S1B and C**). These results indicate that severe skin damage is a cause of class-switching in Dsg3-specific B cells and drives autoimmunity in predisposing individuals to autoimmune disease. Therefore, certain pathogenic conditions may enable Dsg3-specific B cells to execute pathogenic class-switching of immunoglobulin *in vivo*, an indispensable process for pemphigus onset that is prevented under physiological conditions.

### FcγRIIB pathway is downregulated in peripheral blood B cells of pemphigus patients

Although forced Dsg3 exposure promoted class-switching in Dsg3-specific B cells in our mouse model, the relationship between pemphigus onset and preceding severe skin damage remains clinically unclear. To identify novel intrinsic factors associated with B cell pathogenicity in pemphigus patients, we performed genome-wide gene expression analysis of CD19^+^ B cells isolated from pemphigus patients (n = 8) and healthy individuals (n = 9). Gene expression analysis identified 84 of 577 immune-related genes as being differentially expressed between two groups (*P* < 0.05 by edgeR) (**Fig. 4A, and Supplementary Table 1 and 2**). Pathway analysis using these 84 genes identified 57 biological pathways that were affected in pemphigus peripheral B cells (**Fig. S2**). To narrow down the pivotal candidate molecules that are associated with pemphigus pathogenesis, we further analyzed enriched biological functions in patient samples with Integrated Pathway Analysis (IPA) software (**Fig. 4B**). As expected, activation of B lymphocytes was most highly enriched in our dataset and FCGR2B, SEMA4D, and ZAP70, were included among genes altered in the affected biological function. Among them, FcγRIIB is one of the attractive subjects to investigate from the clinical implication, since FcγRIIB function has been not only reported to associate with various autoimmune diseases (22), but more specifically, intravenous immunoglobulin therapy has been effective in treatment of pemphigus vulgaris itself in clinical situation (23–25). Indeed, FcγRIIB signaling was also included in the affected biological pathways (**Fig. S2**). *FCGR2B*, and other downstream molecules of FcγRIIB, including *DOK1*, *BTK*, and *GSK3A*, were downregulated in pemphigus patients compared with healthy individuals (**Fig. 4C**). Meanwhile, *KRAS*, the expression of which is suppressed by FcγRIIB signaling (26), was upregulated in pemphigus patients (**Fig. 4C**). These results suggested that attenuated FcγRIIB pathway is one of the immunological features observed in peripheral blood B cells of pemphigus patients.

**Figure 4.**
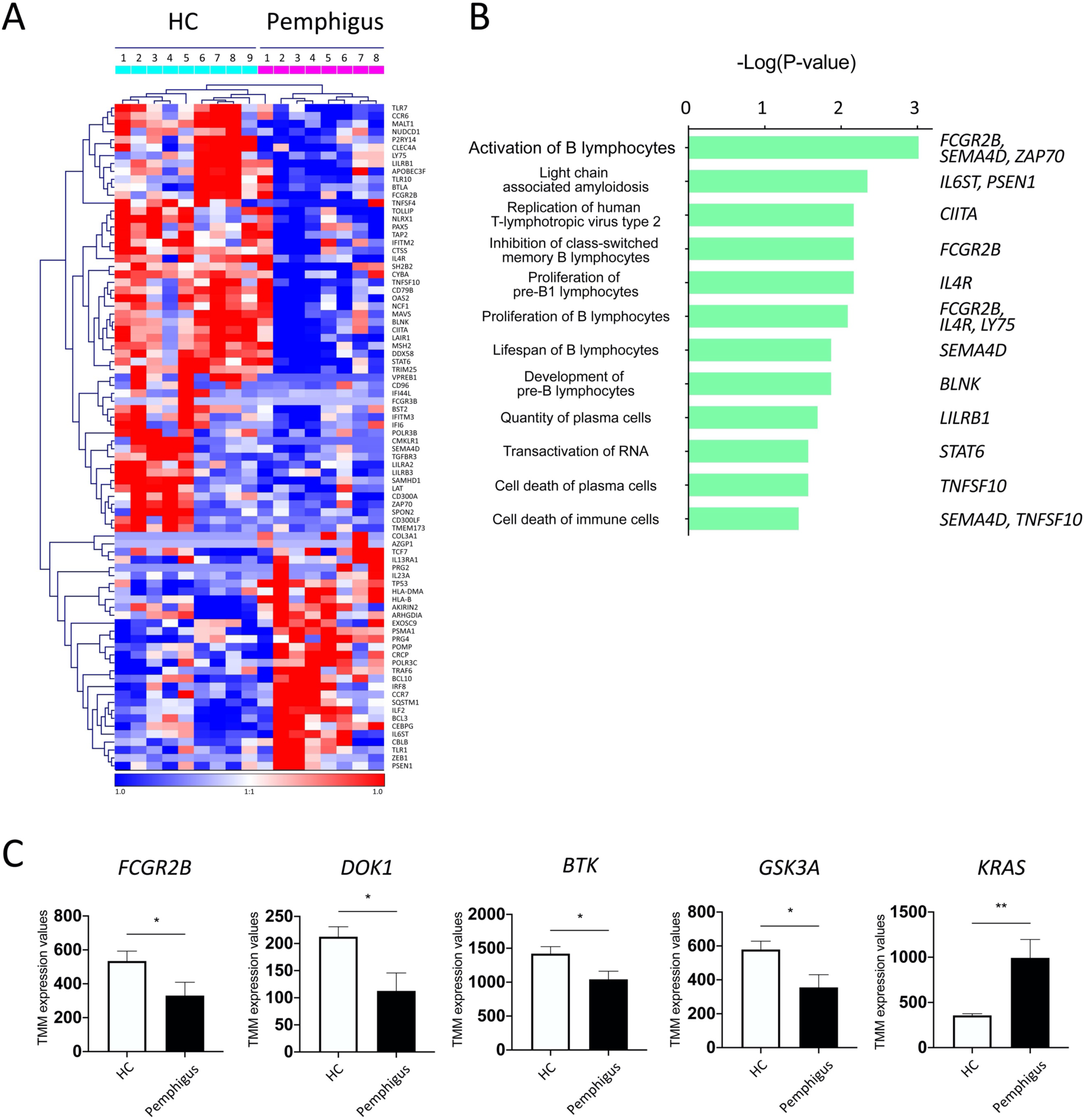
FcγRIIB pathway is downregulated in peripheral blood B cells of pemphigus patients. (**A**) Heatmap of hierarchical clustering of immune-related genes differentially expressed (*P* < 0.05) in peripheral blood B cells from pemphigus patients; (pemphigus vulgaris, n = 6; pemphigus foliaceus, n = 2) and healthy controls (HC; n = 9). See also **Supplementary Table 2** for exact *P* values. (**B**) Enriched biological functions in pemphigus peripheral B cells and genes involved in each function were shown. (**C**) Expression levels (trimmed mean of M values) of FcγRIIB and FcγRIIB pathway-related molecules. \**P* < 0.05, \*\**P* < 0.01; Student’s *t*-test. Data are shown as the mean ± SEM. **Figure 4-Source Data 1** Excel file containing normalized TMM values used for the heatmap shown in **Fig. 4A.** **Figure 4-Source Data 2** Excel file containing the information of related molecules and *P* values of the biological functions analysis with Integrated Pathway Analysis software shown in **Fig. 4B.** **Figure 4-Source Data 3** Excel file containing numerical values of the analyses shown in **Fig. 4B and C**. **Figure 4-figure supplement 2-Source Data 1** Excel file containing the information of related molecules and *P* values of biological pathways analysis with Integrated Pathway Analysis software shown in **Fig. S2**. **Figure 4-figure supplement 2-Source Data 2** Excel file containing numerical values of count data for the biological pathway analysis shown in **Fig. S2**.

### FcγRIIB maintains tolerogenic condition for constraining pathogenic class-switching in AK23H^ki^L^tg^ mouse

Given that attenuated FcγRIIB pathway in B cells was highlighted as one of the characteristic immunological aspects of pemphigus patients via analyzing clinical samples, we asked whether reduced FcγRIIB signaling can authorize Dsg3-reactive B cells to acquire the pathogenicity in the mouse model. To this end, we established an *Fcgr2b*^-/-^-AK23H^ki^L^tg^ mouse line by crossing AK23H^ki^L^tg^ mice with *Fcgr2b*^-/-^ mice on a C57BL/6 genetic background. The body weight of *Fcgr2b*^-/-^-AK23H^ki^L^tg^ mice was significantly lower compared with that of AK23H^ki^L^tg^ mice after 6 weeks of age (**Fig. 5A**). In contrast to AK23H^ki^L^tg^ mice, anti-Dsg3 IgG antibody levels were elevated in *Fcgr2b*^-/-^-AK23H^ki^L^tg^ mice 6 weeks after birth (**Fig. 5B**). However, we observed no differences in total IgG levels between *Fcgr2b^-/-^* and WT mice until 16 weeks after birth (**Fig. 5C**). These results together imply that *Fcgr2b* deficiency itself might not be the single reason that drives early class-switch recombination in *Fcgr2b*^-/-^-AK23H^ki^L^tg^ mice since it did not occur in *Fcgr2b^-/-^ mice*. Rather, autoreactivity of knock-in BCR would presumably be the additional factor that may have accelerated the AK23 BCR class-switch recombination and IgG production in *Fcgr2b*^-/-^-AK23H^ki^L^tg^ mice. Furthermore, *Fcgr2b^+^*^/-^-AK23H^ki^L^tg^ but not AK23H^ki^L^tg^ mice developed PV phenotype as *Fcgr2b*^-/-^-AK23H^ki^L^tg^ mice, including skin erosion and hair loss, as well as acantholytic blister and IgG deposition on keratinocyte cell surfaces (**Fig. 5D**, and data not shown). Additionally, the survival rates of *Fcgr2b^-/-^*-AK23H^ki^L^tg^ and *Fcgr2b^+/-^*-AK23H^ki^L^tg^ mice were significantly lower compared with those of AK23H^ki^L^tg^ mice (**Fig. 5E**).

**Figure 5.**
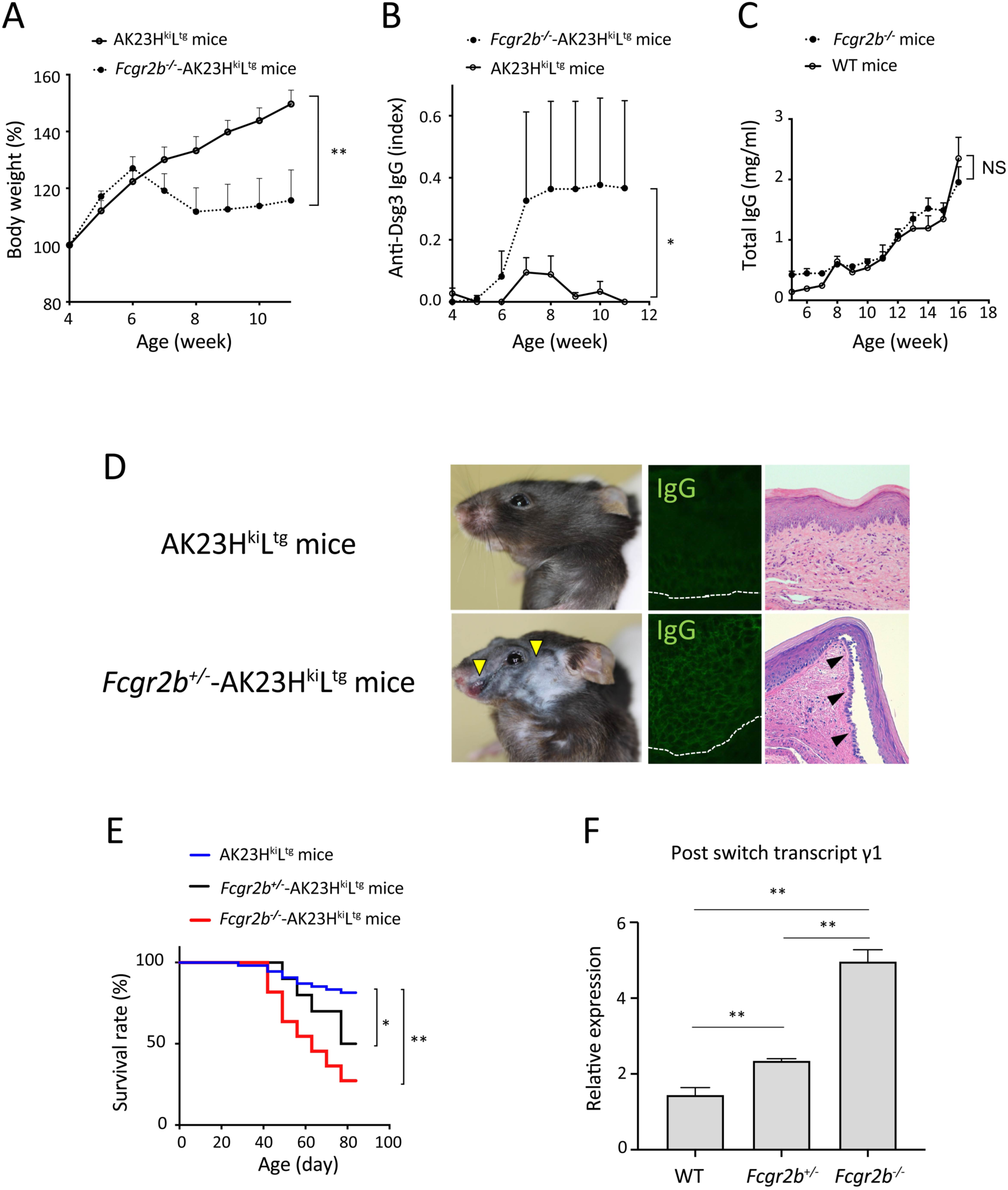
FcγRIIB maintains tolerogenic condition for constraining Dsg3-specific B cell class-switching in AK23H^ki^L^tg^ mice. (**A**) Body weight of *Fcgr2b^-/-^*-AK23H^ki^L^tg^ (closed circle; n = 7) and AK23H^ki^L^tg^ (open circle; n = 6) mice. (**B**) Anti-Dsg3 IgG antibody levels in *Fcgr2b^-/-^*-AK23H^ki^L^tg^ (closed circle; n = 7) and AK23H^ki^L^tg^ (open circle; n = 5) mice were determined by ELISA. (**C**) Non-specific total IgG antibody concentration (mg/mL) in *Fcgr2b^-/-^* (closed circle; n = 6) and WT (open circle; n = 6) mice was determined by ELISA. (**D**) Skin of *Fcgr2b^+/-^*-AK23H^ki^L^tg^ mice. Hair loss and erosion are indicated by yellow arrowheads, IgG depositions are shown in green, and acantholytic blister is indicated by black arrowheads. (**E**) Survival rates of *Fcgr2b^-/-^*-AK23H^ki^L^tg^ (red line; n = 11), *Fcgr2b^+/-^*-AK23H^ki^L^tg^ (black line; n = 10), and AK23H^ki^L^tg^ (blue line; n = 54) mice are assessed in single cohort analysis and shown. (**F**) *Ex vivo* PSTγ1 expression levels in B cells from *Fcgr2b^-/-^* (n = 3), *Fcgr2b^+/-^* (n = 3), and WT (n = 3) mice. \**P* < 0.05, \*\**P* < 0.01. Statistical significance was determined by two-way ANOVA (A, B, C), log-rank test (E), or *t*-test (F). Data are shown as the mean ± SEM. **Figure 5-Source Data 1** Excel file containing numerical values of the analyses shown in **Fig. 5A-C, E and F.**

To elucidate the mechanisms involved in AK23 IgG elevation in the absence of FcγRIIB, we evaluated the class switch efficacy of *Fcgr2b^-/-^*, *Fcgr2b^+/-^*, and WT IgM^+^ B cells *ex vivo*. The expression levels of post-switch transcript (PST) γ1, which is a by-product of class-switch recombination to IgG_1_, were significantly higher in *Fcgr2b^-/-^* and *Fcgr2b^+/-^* B cells compared with those in WT B cells (**Fig. 5F**). Additionally, PSTγ1 levels were significantly higher in *Fcgr2b^-/-^* B cells than *Fcgr2b^+/-^* B cells (**Fig. 5F**). These results imply that FcγRIIB suppresses class-switch recombination in a gene dose-dependent manner, consistent with the differential survival among *Fcgr2b^+/+^*-AK23H^ki^L^tg^, *Fcgr2b^+/-^*-AK23H^ki^L^tg^, and *Fcgr2b^-/-^*-AK23H^ki^L^tg^ mice (**Fig. 5E**). Taken together, FcγRIIB is one of the pivotal factors to prevent pathogenic class-switching of Dsg3-specific B cells and PV phenotype induction in AK23H^ki^L^tg^ mice.

## DISCUSSION

Immunological tolerance is crucial for inhibiting autoreactive lymphocytes and preventing autoimmunity. In fact, interruptions of representative immune tolerance mechanisms, such as deletion, anergy, and receptor editing, lead to autoimmunity (27–29). Given that anti-Dsg3 IgM antibodies are not pathogenic, their class-switching to IgG autoantibodies is essential for pemphigus induction. However, class-switching of anti-Dsg3 IgM antibodies to IgG autoantibodies is absent under physiological conditions in AK23H^ki^L^tg^ mice, even though Dsg3-specific knock-in B cells maintain their ability of class-switching. Our findings imply that a tolerogenic environment is maintained under physiological condition that does not allow immunoglobulin class-switching, even in the presence of corresponding autoantigens. Indeed, pathogenic class-switching was not only forced to occur upon immunization with the recombinant Dsg3 protein, it was also autonomously induced under the reduced FcγRIIB signaling, one of the characteristics of peripheral blood B cells from pemphigus patients. The results of this study demonstrate that inhibition of pathogenic class-switching is a crucial tolerogenic process to avoid pemphigus onset that seems to constantly operate under physiological conditions.

Attenuated FcγRIIB signaling was found to be one of the crucial factors driving pathogenic class-switching in AK23H^ki^L^tg^ mice in this study. FcγRIIB has been reported in many previous studies to inhibit IgG^+^ B cell expansion and antigen presentation by raising the threshold for BCR activation and to induce plasma cell apoptosis etc., mainly related to B cell immune reactions (22, 30). Since class-switching is an immunological event that occurs after BCR activation, it is reasonable to assume that FcγRIIB finely tunes the BCR threshold and modulates immunoglobulin class-switching. Nevertheless, the pivotal role of FcγRIIB in class-switching has not been clearly and experimentally elucidated. In fact, a previous study demonstrated no influence of FcγRIIB deficiency on the proportion of class-switched IgG^+^ B cells in anti-DNA BCR-KI mice (11). In our study, Dsg3-specific IgG^+^ B cells were undetectable in AK23H^ki^L^tg^ mice, whereas the anti-Dsg3 IgG titer spontaneously increased in the *Fcgr2b*^-/-^-AK23H^ki^L^tg^ and, perhaps more importantly, *Fcgr2b*^+/-^-AK23H^ki^L^tg^ mice. Furthermore, FcγRIIB insufficiency in B cells upregulated PSTγ1, an immediate indicator of on-going immunoglobulin class-switching. Together, these results indicate FcγRIIB commitment in the efficiency of immunoglobulin class-switching.

Notably, in contrast to AK23H^ki^L^tg^ mice, AK23H^ki^L^tg^-*Fcgr2b* deficient mice started producing high levels of anti-Dsg3 IgG antibodies 6 weeks after birth (**Fig. 5B**). However, there was no difference in the titers of total IgG antibodies between *Fcgr2b^-/-^* and WT mice until 16 weeks after birth (**Fig. 5C**). The acceleration of class-switching with resultant anti-Dsg3 IgG elevation in the early phase in *Fcgr2b*^-/-^-AK23H^ki^L^tg^ mice can be attributed to the autoreactivity of the knocked-in BCR in the mice. Presumably, immunological stresses, such as expected autoantigen exposure to the knock-in BCR, may have forced class-switching, leading to early pathogenic autoantibody production in AK23H^ki^L^tg^-*Fcgr2b*-deficient mice, but not in AK23H^ki^L^tg^ mice. This could be one of the reasons why FcγRIIB is able to play pathogenic roles in antigen-specific autoimmune diseases, although FcγRIIB itself does not exhibit antigen specificity.

It is impractical to identify the actual immunological stress that drives class-switching in pemphigus patients after disease onset. Immunization with Dsg3 protein and the severe skin damage implemented in this study do not usually occur before pemphigus onset in the clinical situation. On the contrary, it should be noted as shown in this study that *Fcgr2b* haploinsufficiency also accelerated spontaneous class-switching, implying that the reduction in FcγRIIB signaling can trigger spontaneous class-switching. In a previous study, G386C polymorphism in the *FCGR2B* gene promoter, which reduced susceptibility to autoimmune diseases by upregulating FcγRIIB, was less common in pemphigus patients, implying the pathogenic involvement of reduced FCGR2B signaling in pemphigus development (31, 32). More importantly, we demonstrated in this study that the attenuated FcγRIIB pathway identified as one of the phenotypic alterations actually observed in peripheral blood B cells from pemphigus patients was experimentally demonstrated to be crucial for the class-switching of anti-Dsg3 antibody in mice. Although further research is needed to clarify whether FcγRIIb deficiency induces the class-switching in a B cell-intrinsic mechanism, these results implied that reduced FcγRIIB signaling could be one of the predisposing factors in pemphigus development.

Clonal ignorance is another mechanism for immunological tolerance, preventing autoreactive B cell activation (33, 34). Clonal ignorance occurs irrespective of class-switching, since insulin-specific B cells were maintained in a state of clonal ignorance but could exist as IgG^+^ B cells after class-switching (35, 36). In rheumatoid factor (RF) BCR knock-in mice (AM14 sd-Tg mice), B cells remained inactive via clonal ignorance; B cells were activated and underwent class-switching and secreted large amounts of IgG RF antibodies in the MRL/lpr background (37). However, the importance of class-switching in disease pathogenesis could not be investigated in this model, since neither RF IgG nor IgM were pathogenic for phenotype induction. In contrast, PV is a unique autoimmune disease, in which only IgG but not IgM autoantibodies are pathogenic. Therefore, our model was able to specifically focus on class-switching as a tolerance-targeting step, which is completely distinct from previous models. If there is a driving force that constantly induces pathogenic class-switching for harmful tissue damage and there is an opposite force that allows individuals to escape from the harm under physiological conditions, the latter might be immunological tolerance. Our results suggest that inhibition of class-switching is a crucial tolerogenic process to suppress disease progression, at least in pemphigus, a rare disease in which class-switch is indispensable for disease development.

In conclusion, our findings highlight class-switching as the target of the B cell tolerance mechanism, preventing autoimmunity. Further in-depth characterization of the molecular mechanisms downstream of FcγRIIB that is directly connected to class-switching inhibition in autoreactive B cells will be helpful for our understanding of the onset in autoantibody-mediated autoimmune diseases and the development of novel approaches to prevent or treat such conditions.

## MATERIALS AND METHODS

### Mice

AK23 IgM transgenic mice (AK23IgM^tg^) were generated by crossing AK23 IgM H chain transgenic mice with AK23 L chain transgenic mice, both of which have been described previously (17) and were on a C57BL/6 genetic background. C57BL/6-*Rag-2^-/-^* mice (Stock #RAGN12) were purchased from Taconic (Germantown, NY, USA). C57BL/6-*Fcgr2b*^-/-^ mice were purchased from Oriental Bio Service (Kyoto, Japan). C57BL/6 wild-type (WT) mice were purchased from Sankyo Labo Service Corporation (Tokyo, Japan). B6.SJL (CD45.1) WT mice (Stock number 002014) were obtained from The Jackson Laboratory. All mice were maintained under specific pathogen-free conditions at our animal facility. All animal experiments were approved by the Animal Care and Use Committee of the Keio University and RIKEN and were performed in accordance with institutional guidelines.

### Generation of AK23 immunoglobulin knock-in mice

The targeting vector used to create AK23 Ig H chain knock-in mice was constructed by modifying the vector used to generate anti-MOG Ig H chain knock-in mice (38); the vector was kindly provided by Antonio Iglesias (F. Hoffmann-La Roche Ltd, Basel, Switzerland). We replaced the anti-MOG VDJ region with the AK23 VDJ sequence derived from the vector used to generate AK23 IgM transgenic mice (17). The vector contained a neomycin resistance gene cassette and homologous sequences derived from a cosmid vector (**Fig. 1A**). The targeting vector was linearized and electroporated into v6.5 mouse hybrid ES cells (C57BL/6x129/Sv), and 240 neomycin-resistant clones were selected. Five out of 240 clones were determined as transgene-positive by PCR screening. Southern blot analysis was performed using Sac1-digested ES cell DNA and hybridization to a 357 bp DIG-labeled external probe (**Fig. 1A, 1B**). One correctly targeted clone was obtained and the insertion was confirmed by sequencing. The positive ES clone was aggregated with eight-cell *BDF2* embryos and implanted into pseudopregnant hosts. The resulting germline transmitting chimeras were obtained and crossed with C57BL/6 mice to produce AK23 H chain knock-in mice (AK23H^ki^). After backcrossing to C57BL/6 more than 10 times, AK23H^ki^ mice were crossed with AK23 L chain transgenic mice (17) for generating AK23 immunoglobulin knock-in mice (AK23H^ki^L^tg^).

### Bone marrow transfer experiment

CD45.2^+^ WT and CD45.2^+^ *Dsg3^-/-^* mice were lethally irradiated. The minimum lethal doses of CD45.2^+^ WT and CD45.2^+^ *Dsg3^-/-^* mice were 9–9.5 Gy and 7.5–8 Gy, respectively. After irradiation, they received bone-marrow cells from AK23H^ki^L^tg^-*Rag2^-/-^* mice (CD45.1^+^).

### *In vitro* class-switching and adoptive transfer

B cells were purified from splenocytes by positive selection using magnetic beads-conjugated monoclonal anti-mouse B220 antibody (clone: RA3-6B2, Miltenyi Biotec, Bergisch Gladbach, Germany). Isolated B cells (5 × 10^5^/mL) were incubated with IL-4 (20 ng/mL) and LPS (10 μg/mL) for 96 h. Then, 8 × 10^6^ B cells were adoptively transferred into *Rag2^-/-^* mice via the tail vein.

### Direct immunofluorescence

Palate tissues were embedded in OCT compound (Tissue-Tek; Sakura Finetechnical, Tokyo, Japan), frozen immediately at -140°C, and cut into 6-μm-thick sections. Sections were incubated with Alexa 488 anti-mouse IgG_1_ Ab (Invitrogen) for 1 h at room temperature, washed in PBS, cover-slipped using Mowiol (Calbiochem, Darmstadt, Germany), and imaged using a fluorescence microscope.

### ELISA

Serum anti-Dsg3 Ig titers were determined by ELISA using plates coated with 5 μg/mL (**Fig. 1, 4**) or 3 μg/mL (**Fig. 6**) rDsg3 protein as described previously (15). Each serum sample was diluted 1:200, 1:500, and 1:1000. Sera obtained from PV model mice (15) and culture supernatant of AK23 hybridoma cells (16) were used as positive control for anti-Dsg3 IgG. AK23IgM^tg^ mice (17) were used as positive controls for anti-Dsg3 IgM. The index value for anti-Dsg3 IgG was calculated as previously described (14). Total IgG antibody concentrations were determined using the IgG Mouse ELISA kit (Abcam, Cambridge, UK) according to the manufacturer’s instructions.

### Flow cytometry

Single-cell suspensions were prepared from mouse peripheral blood, spleens, and lymph nodes. Cells were stained with the following antibody conjugates: biotin-conjugated anti-IgM^a^ (DS-1; BD Biosciences), anti-IgG_1_ (B68-2; BD Biosciences), anti-BP-1 (6C3; Biolegend), and anti-IgM (RMM-1; Biolegend); FITC-conjugated anti-IgM^a^ (DS-1; BD Biosciences) and anti-E-tag (Bethyl Laboratories, Montgomery, TX, USA); PE-conjugated anti-IgM^b^ (AF-6-78; BD Biosciences), anti-CD23 (B3B4; BD Biosciences), anti-IgM (AF6-78; BD Biosciences), and streptavidin; APC-conjugated anti-B220 (RA3-6B2; BD Biosciences), anti-CD24 (M1/69; Biolegend), anti-IgD (11-26c.2a; Biolegend), and streptavidin; APC-Cy7-conjugated anti-CD21 (7E9; Biolegend); PE/Cy7-conjugated anti-IgM (RMM-1; Biolegend); PB-conjugated anti-B220 (RA3-6B2; Biolegend), and streptavidin.

To determine Dsg3-binding to B cells, we purified B cells (**Fig. 1D**) by positive selection using magnetic beads (MACS, Miltenyi Biotec) conjugated with a monoclonal anti-mouse B220 antibody (clone: RA3-6B2; Miltenyi Biotec) according to the manufacturer’s instructions. Subsequently, cells were incubated with 10 μg of rDsg3-E-His in 100 μL of RPMI on ice for 1 h and stained with anti-E-tag-FITC (Bethyl Laboratories). After excluding dead cells by staining with 7AAD, cells were analyzed with a FACSCanto II flow cytometer (BD Biosciences), and the resulting data were analyzed using FlowJo software (Tree Star, Ashland, OR, USA).

### Calcium mobilization

Splenocytes were stained with anti-CD45.1-PE/Cy7 and anti-B220-APC (Biolegend, San Diego, CA, USA) at 4°C for 30 min in PBS . Then, cells were incubated in HBSS containing 10 µM Fluo-4 (Molecular Probes, Eugene, OR, USA) at room temperature for 30 min. Following stimulation with 20 µg/ml anti-IgM antibody, calcium mobilization was measured on a FACSCanto II flow cytometer (BD Biosciences, San Jose, CA, USA).

### Immunization with rDsg3

A recombinant baculoprotein of mouse Dsg3 that included the extracellular domain of mouse Dsg3, an E-tag, and a His-tag was used for immunization as described previously (15). Mice were primed by subcutaneous injection of purified rDsg3 (5 μg) in complete Freund’s adjuvant. Subsequently, mice were boosted twice with rDsg3 in incomplete Freund’s adjuvant, following by two intraperitoneal injections of rDsg3 without adjuvant, once per week.

### Hapten-induced dermatitis

A total of 80–100 μL of 0.5% 1-fluoro-2,4-dinitrobenzene (DNFB; Nacalai Tesque, Kyoto, Japan) in acetone/olive oil (3:1) was applied to the shaved abdomens or backs of mice weekly up to 4-7 times, depending on the experiment.

### Vaccinia virus infection

Vaccinia virus strain VV-WR was a kind gift from Masayuki Saijo (National Institute of Infectious Diseases, Department of Virology I, Japan). Mice were infected with VV-WR by skin scarification. Briefly, under anesthesia, abdomens or backs of mice were shaved and scratched with a 27G needle. VV-WR (1 × 10^8^ PFU/mL, 150–200 μL/mouse) was applied to the scarred skin areas every 2 or 3 weeks.

### RNA sequencing and transcriptomic analysis

The study protocols were reviewed and approved by the institutional review board of the Keio University School of Medicine and RIKEN, and were conducted following the principles established by the Declaration of Helsinki. Written informed consent was obtained from all patients. B lymphocytes in peripheral blood from 8 pemphigus patients (six pemphigus vulgaris and two pemphigus foliaceus patients), and nine healthy donors, were identified as CD3^-^CD19^+^ lymphocytes and sorted with a FACSAria III flow cytometer (BD Biosciences). Total RNA was extracted from the sorted lymphocytes using TRIzol reagent (Invitrogen, Carlsbad, CA, USA). The detailed clinical information of the patients is shown in **Supplementary Table 1**. RNA-seq libraries were prepared using the NEBNext Ultra RNA Library Prep Kit for Illumina (New England Biolabs, Ipswich, MA, USA) according to manufacturer’s instructions and sequenced using HiSeq2500 (Illumina, on a 50-base single-end read mode). The sequence reads were mapped to the hg37 reference genome (UCSC) using TopHat2 version 2.0.8 and botwie2 version 2.1.0 with default parameters and gene annotation was provided by NCBI RefSeq.

The transcript abundances were estimated using Cufflinks (version 2.1.1). The negative binomial model-based method edgeR (3.10.0) was used for differential expression analysis from raw count data. Normalized trimmed means of M-values (TMM) were visualized in a heatmap.

Affected biological pathways and functions (*P* value < 0.05) in pemphigus peripheral B cells were identified through analyses by Canonical Pathway and Diseases and Biological Functions provided in Ingenuity Pathway Analysis software (IPA, Qiagen, Germantown, MD) (39).

### Analysis on the RNA expression levels of PST γ1 in IgM^+^ cells ex *vivo*

Splenic IgM^+^ cells were isolated by positive selection using magnetic beads (MACS, Miltenyi Biotec). Total RNA of the IgM^+^ cell was extracted using the RNeasy mini kit (Qiagen), and cDNA synthesis was performed using the TaqMan Reverse Transcription Reagent (Applied Biosystems, Foster City, CA, USA). Reactions were prepared using the Universal SYBR Select Master Mix, and quantitative PCR was performed using a StepOnePlus Real-Time PCR System (Applied Biosystems) and the following primers: HPRT forward, 5’-AGCCTAAGATGAGCGCAAGT-3’; HPRT reverse, 5’-TTACTAGGCAGATGGCCACA-3’; post-switch transcript γ1 forward, 5’-ACCTGGGAATGTATGGTTGTGGCTT-3’; post-switch transcript γ1 reverse, 5’-ATGGAGTTAGTTTGGGCAGCA-3’.

### Statistical Analysis

All data are shown as mean ± S.E.M. Data were analyzed by Student’s *t*-test unless specified otherwise. Significance was set at a *P* value of less than 0.05.

## ACKNOWLEDGMENTS

We are grateful to Drs. Antonio Iglesias and Masayuki Saijo for providing the anti-MOG Ig H chain knock-in construct and the vaccinia virus, respectively; Drs. Sayuri Chiba and Aiko Shiohama for their guidance in generating the DNA constructs; Dr. Hidehiro Fukuyama for fruitful scientific advice; Ms. Kyoko Hidaka for animal care and genotyping of the mice; Ms. Minae Suzuki and Ms. Hiroyo Koike for the preparation of cryosections; and Ms. Mariko Okajima for laboratory managements.

## Supplementary Materials

**Figure S1.**
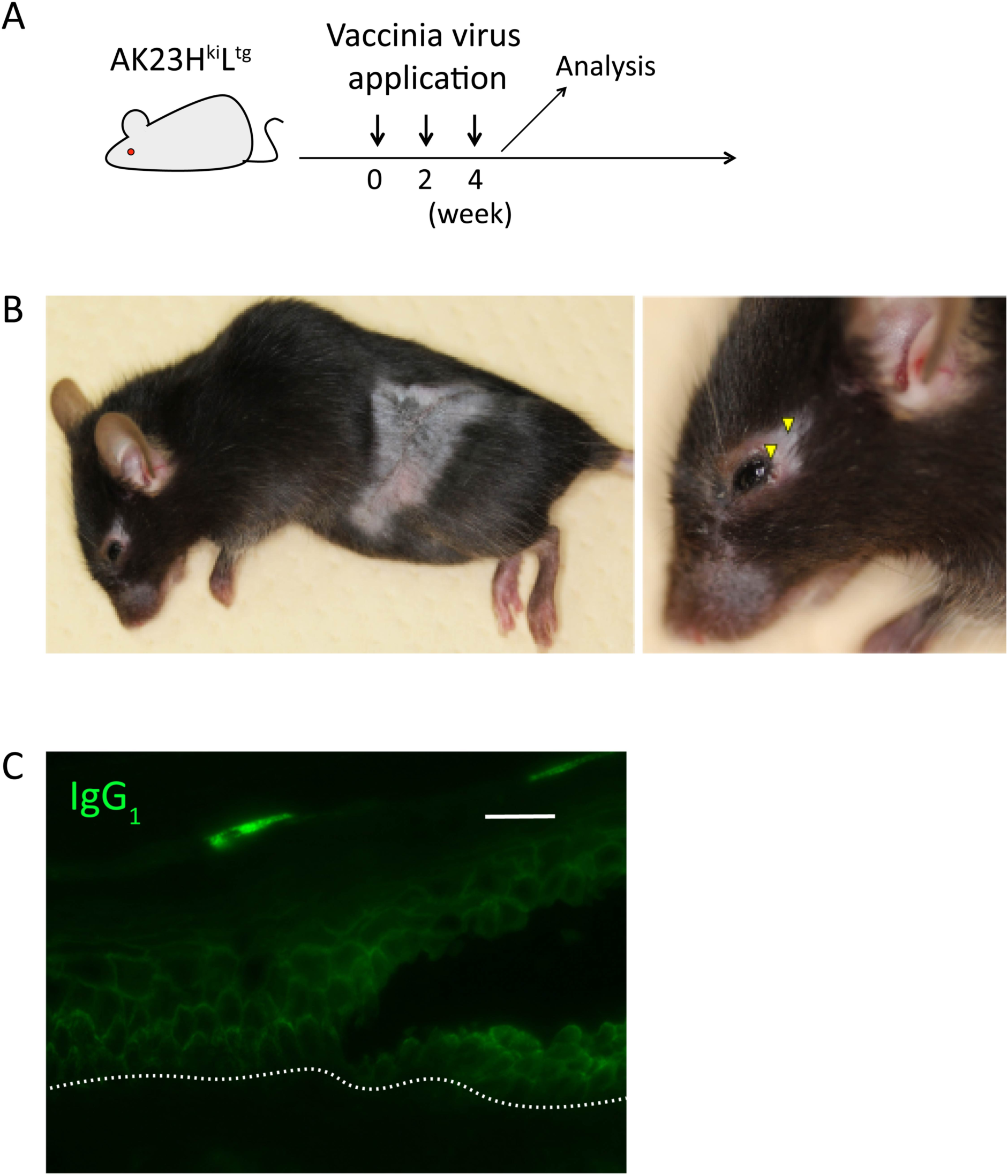
Forced class-switching in Dsg3-specific BCR knock-in B cells by vaccinia virus inoculation induces PV phenotype *in vivo*. (**A**) Schematic representation of vaccinia virus inoculation in AK23H^ki^L^tg^ mice (n = 2) to force class-switching *in vivo*. (**B**) Skin phenotype of AK23H^ki^L^tg^ mice after vaccinia virus inoculation. Hair loss and erosion in the periorbital area are indicated by yellow arrowheads. (**C**) IgG_1_ depositions are shown in green. Data are representative of two independent experiments. Scale bar, 20 μm.

**Figure S2.**
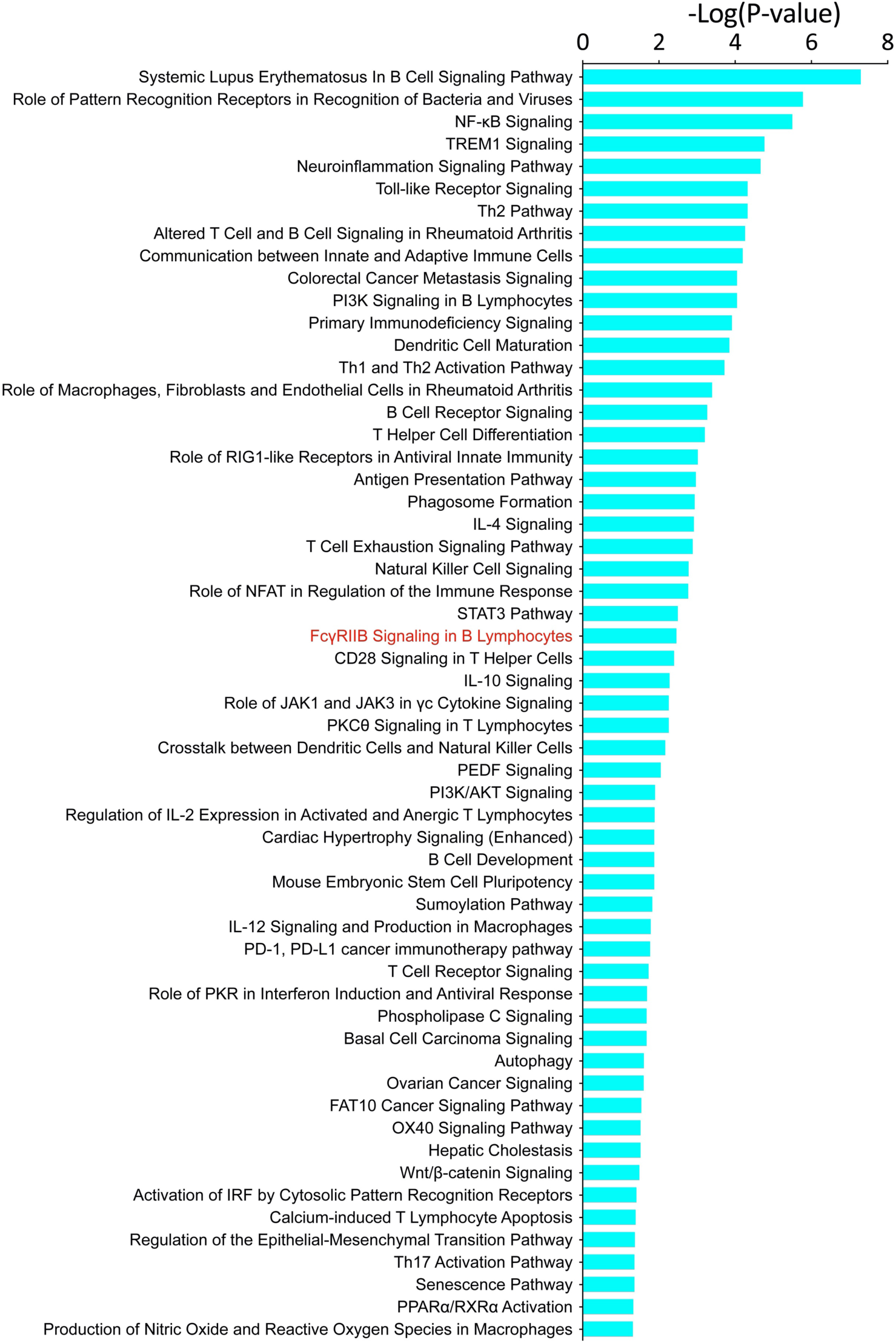
Affected biological pathways in peripheral blood B cells of pemphigus patients identified by canonical pathway analysis of IPA. The results of pathway analysis, using the genes that are both annotated as immunity and detected as *P* value < 0.05 by EdgeR.

**Figure S3.**
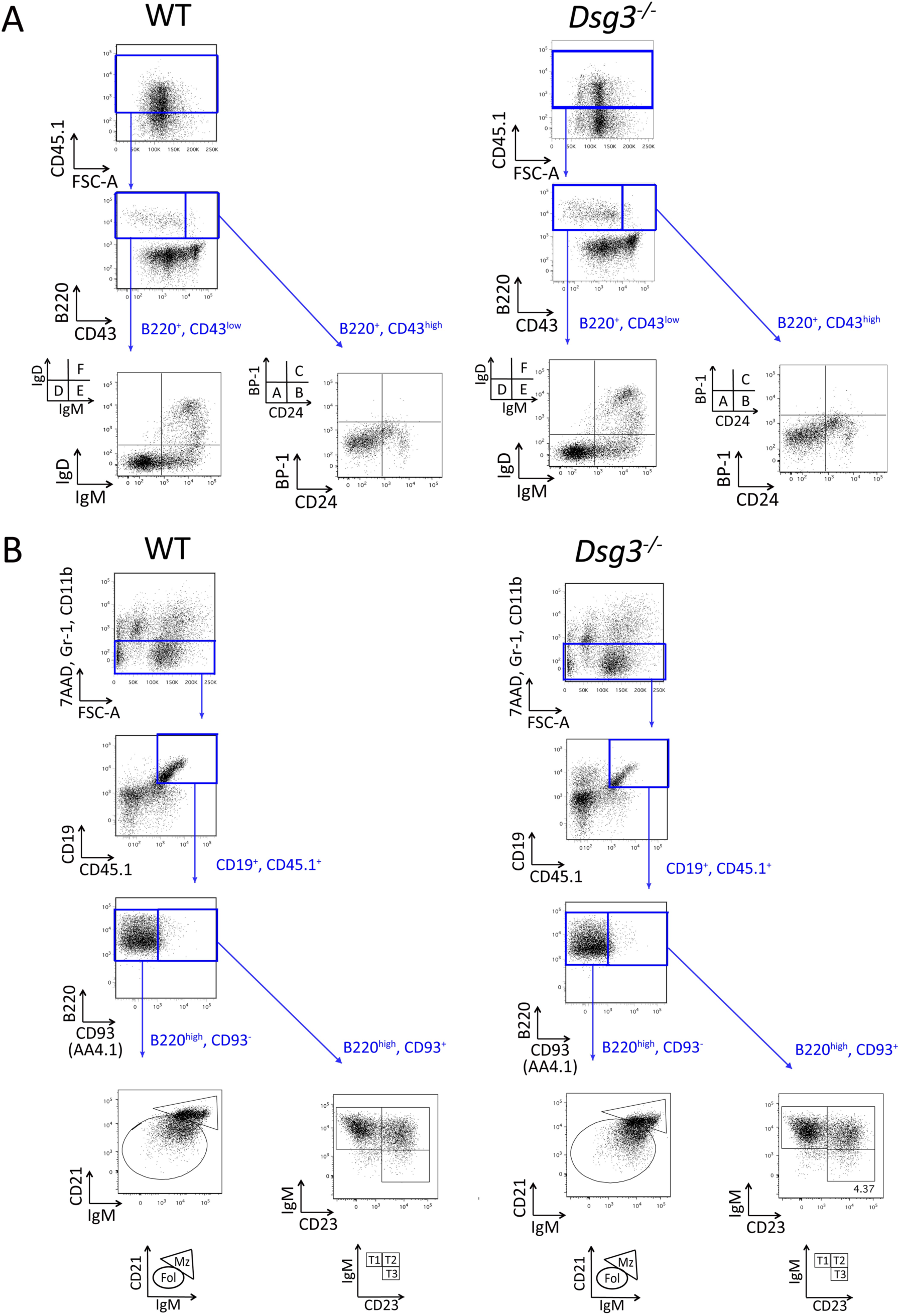
Gating strategy to assess AK23H^ki^L^tg^ B cell development. (**A**) After BMT shown in Fig. 2A, Hardy fraction A–C of donor-derived B cells were detected after gating on CD45.1^+^B220^+^CD43^high^ cells in WT and *Dsg3^-/-^* recipients. Hardy fraction D–F of donor-derived B cells were detected after gating on CD45.1^+^B220^+^CD43^low^ cells in WT and *Dsg3^-/-^* recipients. Representative gating strategies are shown. (**B**) Donor-derived B cells at transitional stages (T1, T2, and T3) were evaluated after gating on CD45.1^+^Gr-1^-^CD11b^-^ CD19^+^B220^high^CD93^+^ cells in WT and *Dsg3^-/-^* recipients. Follicular (Fol) and marginal zone (Mz) donor-derived B cells were evaluated after gating for CD45.1^+^Gr-1^-^CD11b^-^ CD19^+^B220^high^CD93^-^ cells in WT and *Dsg3^-/-^* recipients. 7AAD^+^Gr-1^+^CD11b^+^ cells were excluded. Representative gating strategies are shown.

**Table S1.** Clinical information of the pemphigus patients

| Patient no. | Age (years) | Sex | Pemphigus subtype | IgG antibody titer (CLEIA; U/mL) |  | Titer of IgG antibody (ELISA; Index) |  | PDAI |
| --- | --- | --- | --- | --- | --- | --- | --- | --- |
|  |  |  |  | Anti-Dsg1 | Anti-Dsg3 | Anti-Dsg1 | Anti-Dsg3 |  |
| 1 | 60 | M | mPV | < 3.0 | 561 | NA | NA | 25 |
| 2 | 50 | M | PF | 318 | < 3.0 | NA | NA | 6 |
| 3 | 49 | M | mPV | NA | 191 | NA | NA | 9 |
| 4 | 60 | F | mcPV | 41 | 139 | 220.1 | 271.3 | 25 |
| 5 | 75 | F | mPV | < 3.0 | 69.9 | NA | NA | 50 |
| 6 | 70 | F | PF | 395 | NA | NA | NA | NA |
| 7 | 44 | F | mcPV | 1820 | 7.6 | 586.2 | 50.1 | 35 |
| 8 | 52 | F | mcPV | NA | 730 | NA | NA | 7 |

Abbreviations: M, male; F, female; PV, pemphigus vulgaris; PF, pemphigus foliaceus; mPV, mucosal-dominant type PV; cPV, cutaneous type PV; mcPV, mucocutaneous type PV; CLEIA, chemiluminescent enzyme immunoassay; ELISA, enzyme-linked immunosorbent assay; Dsg, desmoglein; PDAI, pemphigus disease area index; NA, not available

**Table S2.**
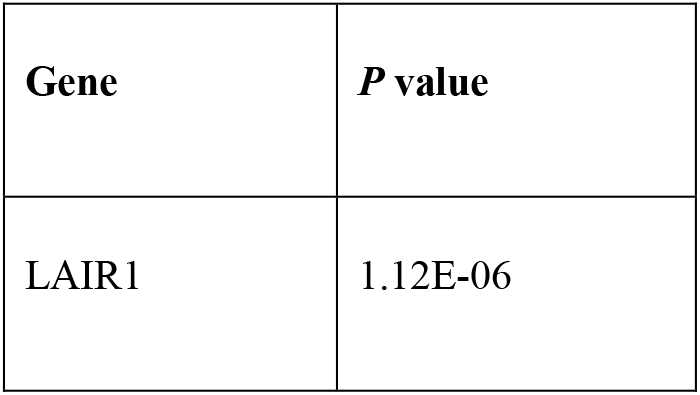
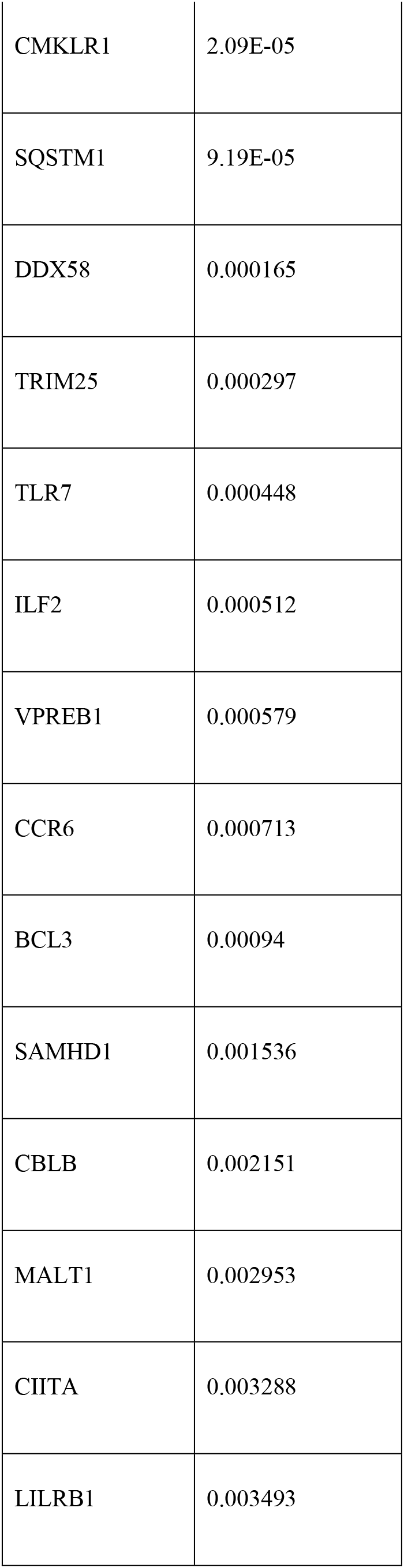
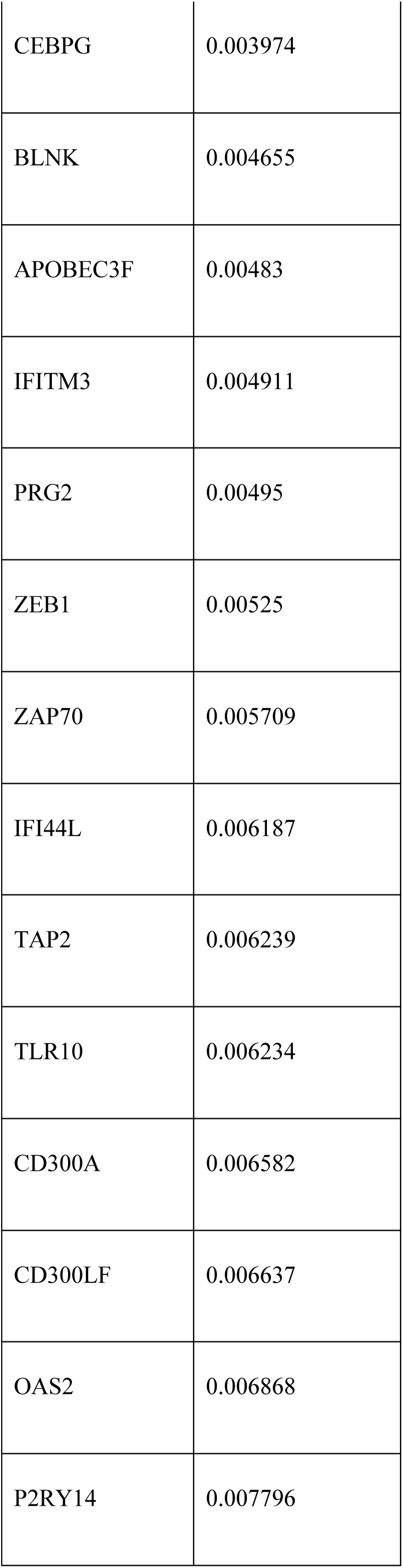
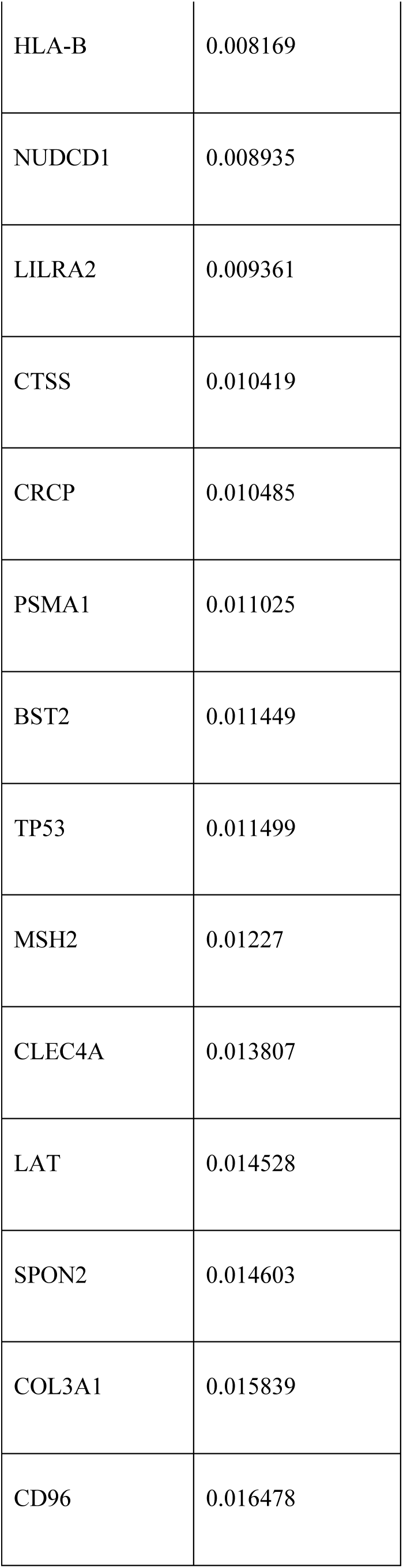
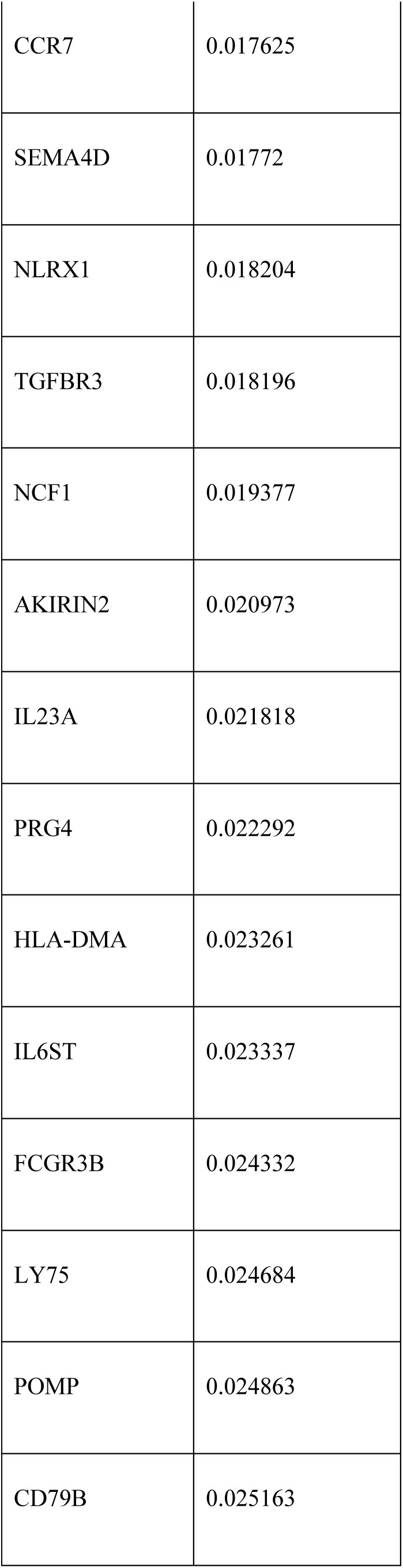
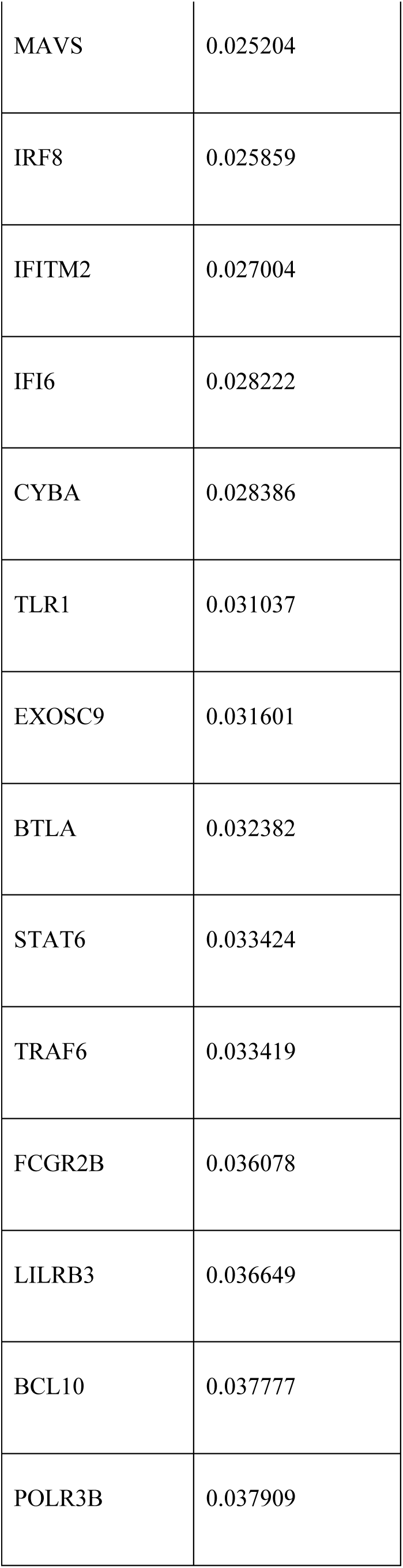
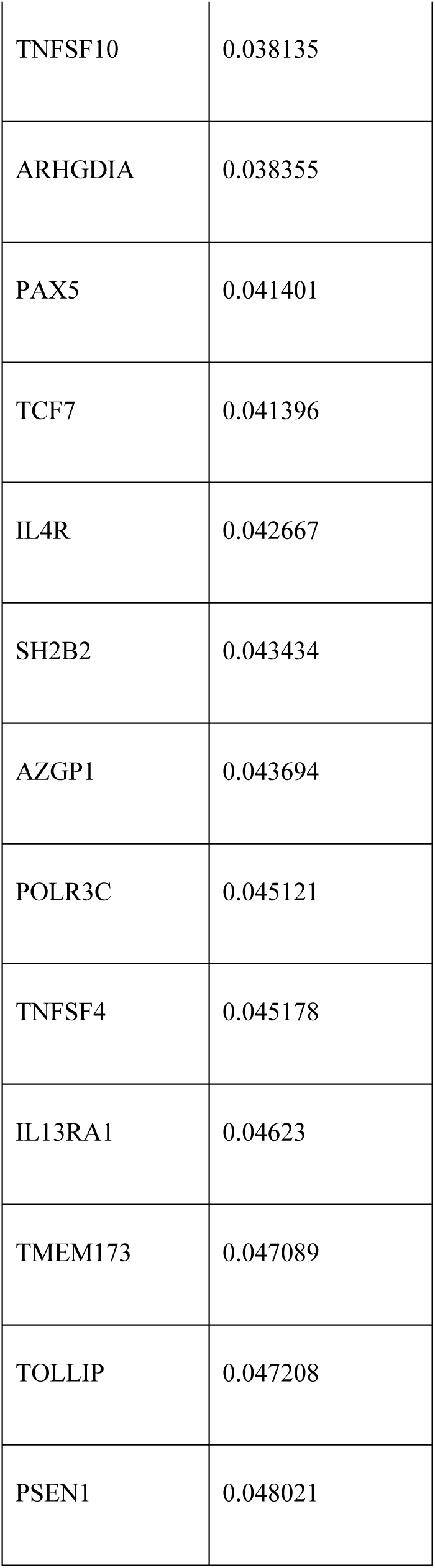
*P* values of genes by edgeR

